# Transcriptional activity of TRF2 is telomere length dependent

**DOI:** 10.1101/056481

**Authors:** Ananda Kishore Mukherjee, Shalu Sharma, Parashar Dhapola, Dhurjhoti Saha, Tabish Hussain, Sumitabho Deb Roy, Gunjan Purohit, Anirban Kar, Ankita Singh, Suman Sengupta, Vivek Srivastava, Manish Kumar, Sagar Sengupta, Shantanu Chowdhury

## Abstract

TRF2 is a telomere repeat binding factor crucial for telomere maintenance and genome stability. An emerging non-conventional role of TRF2 is as a transcriptional regulator through extra-telomeric bindings. Herein we report that increase in telomere length leads to sequestration of TRF2 at the telomeres leading to reduced extra-telomeric TRF2 occupancy genome wide. Decrease in TRF2 occupancy was found on multiple gene promoters in cells with elongated telomeres, including the cell cycle regulator kinase-*p21*. We found that TRF2 is a transcriptional repressor of *p21*, and, interestingly, TRF2-mediated regulatory control of *p21* is telomere length dependent.

---

TRF2,as part of the shelterin complex, confers stability to telomeres^1–4^. Primarily studied as a telomeric protein, it was only recently that genome wide TRF2-binding sites were detected outside the telomeres-many such sites were interstitial telomeric-repeat sequences (ITS)^5^. Genomic binding supports extra-telomeric TRF2 functions, including recently noted transcription^6^ and DNA repair^7^-however, underlying mechanisms are not well-understood. Moreover, recent evidence demonstrates TRF2's potential as a transcription factor in regulation of *PDGFRβ's* expression^6^. These findings bring to light an interesting partitioning of TRF2's genomic occupancy into two types of regions: the extra-telomeric regions of more or less constant cumulative length and telomeric regions of highly variable lengths. This unique genomic distribution of TRF2 led us to hypothesize a ‘sequestration model’ where longer telomeres may alter extra-telomeric occupancy of TRF2, and thus affect TRF2-mediated transcription.

On performing ChlP-sequencing of endogenous TRF2 in fibrosarcoma cells HT1080, we found that about 40% of TRF2 ChlP-sequencing reads were from telomere regions. We also noted that 9.39% of 15.79 million aligned reads mapped to interstitial TTAGGG repeats and there were a total of 20347 peaks (called using MACS) common between two replicates. On finding a large number (3635) of TRF2 peaks within 10 kb of transcription start sites (TSS) we analyzed the TRF2-dependent transcriptome. Following stable over-expression of TRF2 in HT1080 cells, we performed RNA sequencing and found that 171 of up-regulated genes and 503 of down-regulated genes were significantly enriched for sequence reads contributing to TRF2 peaks (Figure 1A). A closer inspection of randomly chosen loci with TRF2 peaks showed consistent TRF2 ChIP signal between replicates (Figure 1B); these and several genomic regions with TRF2 peaks were validated using ChlP-PCR (Figure 1C). A significant number of genes that were differentially expressed and harbored TRF2 peaks were involved in signaling pathways that have implications in cancer and senescence (Supplementary Figure 1B).

**Figure 1.**
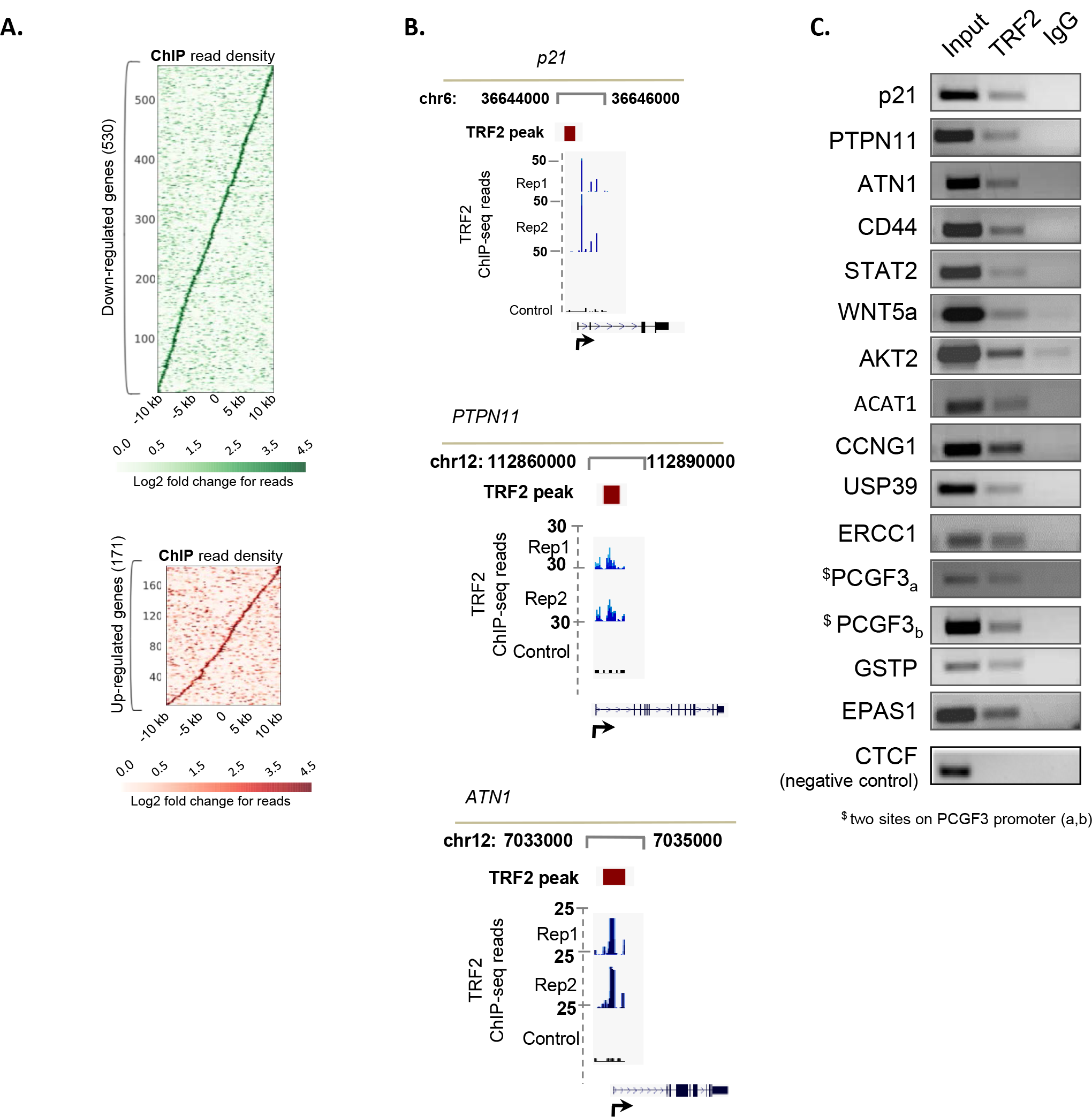
TRF2 has a large number of extra-telomeric binding sites across the genome. **A. Genome wide TRF2 peaks detected:** ChlP-seq read intensity (normalized to input sample) of TRF2 binding within promoters of up or down regulated genes obtained from RNA seq analysis. Here genes have been sorted by the distance of maximum intensity of TRF2 signal on a promoter from TSS. **B. TRF2 binds promoter G4-motifs:** *p21,PTPN11* and *ATN1* promoters showing TRF2 peaks along with raw signal as obtained in two different ChIP-Seq replicates and input control. **C. Validation of TRF2 occupancy at interstitial sites** TRF2 occupancy was observed in randomly selected gene promoters. CTCF promoter, which does not contain TRF2 binding peak within +/– 5 kb to TSS was used as a negative control locus.

G-quadruplexes are secondary DNA structures that are profusely abundant in the telomeric regions and it is known that these structures are crucial for recruitment ^12^ of TRF2 on the telomeres. Computationally, these structures are frequently studied as sequence patterns that can adopt G4-motif or potential G4-motif (PG4-motif in following text to denote computationally predicted G4-motifs) comprising four runs of three guanines (G3) with three loops of up to 15 bases (denoted by the notation (G_3_L_1-15_). PG4-motifs,irrespective of loop length were enriched within TRF2 peaks; for instance, on considering structures with loops of 1-7 bases, 2831 PG4-motifs were present within 1568 (7.7%) of the20347 TRF2 peaks (p<0.01; Figure 2A). Conversely, TRF2 peaks were also enriched withinPG4-motif sequences (p<0.01, Figure 2B). A vast majority (∼95% of 701) of differentially expressed genes harbored at least one potential quadruplex (PG4) motif (G_3_L_1-7_) within 10 kb of TSS. We also found that PG4 motifs were more proximal to promoter-TRF2 peaks (3635 peaks within 10 kb of TSS of 3310 genes; (Figure 2C) than randomly expected. Additional evidence supporting colocalization of extra-telomeric TRF2 and G4-motifs was obtained by immunofluorescence using the recently reported G4-specific antibody 1H6^13^along with TRF2 antibody. We found that the co-localization of TRF2 and G4 signal remains within a significant range of 60-80% across three different cell lines (Figure 2D). The synthetic pyridine based derivative 360A (2,6-N,N-methyl-quinolinio-3-yl)-pyridine dicarboxamide) was shown to interact highly selectively *in vitro* with G4s and also had intracellular affinity for G4-motifs^14^. Treatment of three different cell lines with 360A led to a consistent decrease in nuclear TRF2 signal intensity (>30% (0.5 uM 360A) or >70% (1uM 360A) reduction over untreated cells) (Figure 2E). We further tested the role of G4 DNA in TRF2 occupancy taking a candidate gene approach using genes that featured TRF2 promoter-peak in ChlP-Seq data. We observed that these genes upon 360A treatment show reduced TRF2 occupancy (Figure 2F). We also tested the expression of these genes upon 360A treatment and found an altered gene expression in all the cases (Figure 2G).

**Figure 2.**
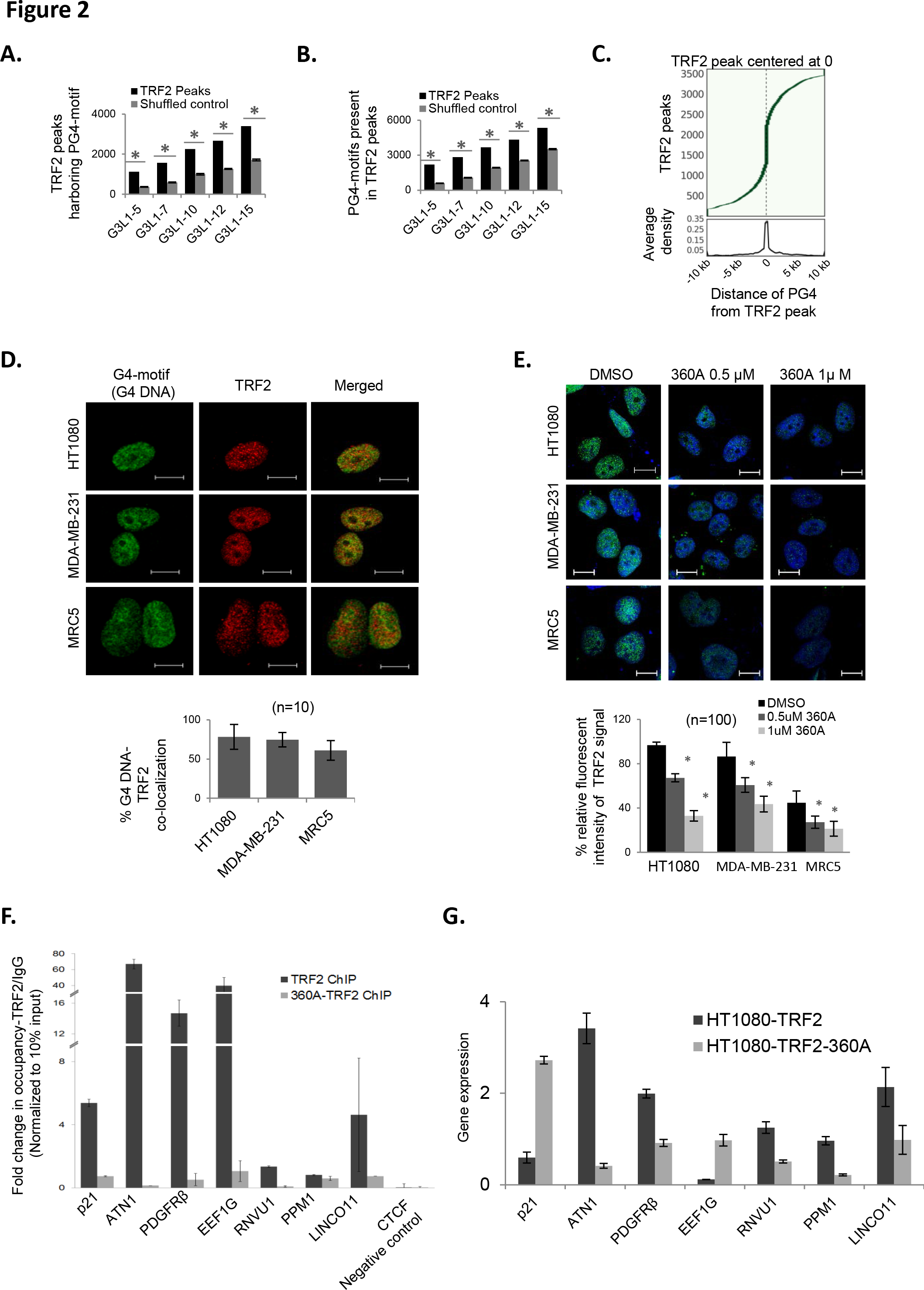
TRF2 binding to multiple extra-telomeric sites is Quadruplex dependent. **Significant number TRF2 peaks show close proximity to putative G4 motifs**. **(A)** A significant number of TRF2 peaks have enriched presence of PG4 motifs (with respect to G4 count following shuffling of respective sequences 100 times; p<0.05). **(B)** G4 motifs sequences were also found to be enriched within TRF2 peaks with respect to counts when each region was shuffled 100 times keeping GC content identical; *: p< 0.05 (Fisher's exact test) **(C)** A large number of TRF2 peaks had putative G4 motifs within a 10 kb window with a significant enrichment for G4 motifs within 5 kb of TSS. **TRF2 associates with G4-motifs. (D)** G4-motif specific antibody 1H6 (Henderson *et al.,* 2014) co-localize with TRF2 in cancer cells, HT1080 and MDA-MB-231 and normal fibroblasts MRC5 (scale bar=10 μm). Percent G4-TRF2 co-localization is given in adjacent histogram; average of readings from ten independent frames. **(E)** Reduced TRF2 signal observed on pre-treatment with G4-specific ligand 360A in HT1080, MDA-MB-231 and MRC5 cells (scale bar=10 μm). Area outlined in HT1080 panel zoomed to show the reduction in TRF2 signal (white arrow) with 360A treatment. Quantitative analysis of TRF2 signal intensity in 360A-treated or control (DMSO-treated) cells given in histogram; data represents quantitation of 100 independent cells; *: p< 0.05 (unpaired Student's t-Test). **G-quadruplex ligand 360A alters TRF2 occupancy at multiple gene promoters leading to changes in expression of the genes(F-G).** Upon 360A treatment a reduction in TRF2 occupancy **(F)** as well as change in expression **(G)**was observed for multiple genes selected from ChlP-Seq data in HT1080 cells.

Together ours' and recent data from other groups^5,6,11^ suggest that telomeric localization of TRF2 does not engage all the TRF2 molecules in the nucleus. Therefore, TRF2 is available for extra-telomeric binding. In the backdrop of this knowledge, we asked if telomeres play any role in either release or sequestration of TRF2. We therefore, wished to check if a longer telomere would be able to sequester more TRF2 molecules and hence decrease the extra-telomeric binding of TRF2 to certain gene promoters. HT1080 isogenic cell line, HT1080-Super Telomerase (HT1080-ST) cells with enhanced telomeres were obtained as a gift from Lingner's lab. These cells constitutively express both *hTERT* and *hTERC* through genomic insertion of both the cDNAs downstream of a CMV promoter^15^ and have an enhanced telomere length compared to HT1080 wild type cells. We re-characterized the cells and found that the cells have approximately 2.5-fold higher telomere length compared to wild type H1080 cells (Figure 3A, Supplementary Figure 2A). Telomere occupancy of TRF2 increased with increase in telomere length in HT1080-ST cells (Figure 3B), which indicates that moreTRF2 molecules are sequestered by longer telomeres. Chromatin bound TRF2 did not show any detectable difference in HT1080-ST cells and HT1080 cells, and nucleoplasm fraction (unbound TRF2 in the nucleus) gave much weaker signals relative to chromatin bound TRF2 for the same amount of protein lysate (Supplementary Figure 2C). This suggested that most of the TRF2 molecules in the nucleus are in a chromatin bound form. Interestingly, when we compared the ChlP-seq data of HT1080-ST with HT1080, we found that there was an overall decrease in extra-telomeric TRF2 occupancy in ST cells (See methods) (Figure 3C). We validated the trend that was observed in the ChlP-seq data using 10 candidate promoters that had shown strong occupancy in the endogenous TRF2 ChlP-seq in HT1080 cells and seven such promoters showed reduced TRF2 occupancy in ST cells with elongated telomeres (data for the seven promoter sites have been shown in Figure 3D). Next, we were interested to know if this change in TRF2 occupancy has any functional consequence on the expression of the candidate genes. Five out of the seven sites showed differential expression (Figure 3E). It was found recently that TRF2 is an activator of *PDGFR β,* that is, loss of TRF2 occupancy at the *PDGFRβ* promoter lead to a down-regulation of the gene^6^. Among the genes that showed differential expression, *p21* is a well-known gene with crucial implications in cell cycle regulation and cellular phenomena ^8,9^ like senescence and apoptosis. We chose to focus on *p21* for further experiments. We found that transient knockdown of TRF2 (which is expected to decrease TRF2 occupancy on the *p21* promoter) led to increase in *p21* promoter activity and *p21* protein levels (Figure 3F), indicating transcriptional repression of *p21* by TRF2. Furthermore, in line with decreased occupancy of TRF2 at the *p21* promoter (Figure 3D) in ST cells relative to HT cells, we found that *p21* promoter activity, mRNA expression and protein level was enhanced in HT1080-ST cells relative to HT1080 cells (Figure 3G).

**Figure 3.**
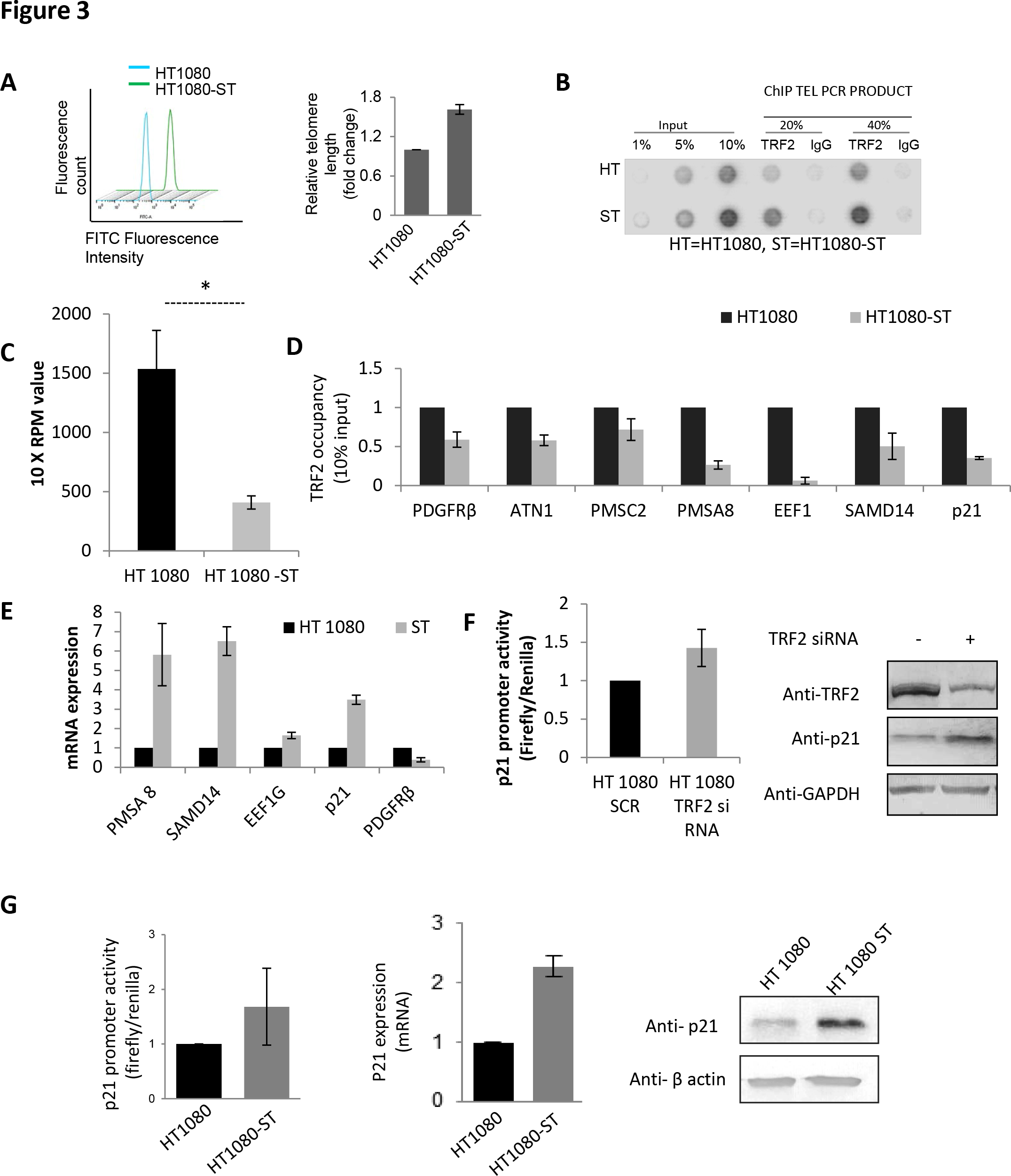
Reduced TRF2 occupancy at several gene promoters in cells with elongated telomeres lead to transcriptional modulation. **A.Telomere length elongation in HT 1080-Super Telomerase (ST) cells (A)**Telomere elongation in oligo-fed HT 1080-ST cells was confirmed by flow-FISH using FITC-tagged Telomere specific probe. **B. Telomere elongation in HT 1080-ST cells show increased telomere occupancy of TRF2 in comparison to HT 1080 cells (B)** ChIP with TRF2 antibody (or isotypic control) followed by PCR using telomere-specific primer in HT 1080 and HT 1080-STcells showed that increase in telomere length resulted in increase in TRF2 occupancy at telomeres in ST cells **C. Telomere elongated HT 1080-ST cells have decreased TRF2 signal in interstitial regions**. **(C)** 1226 genomic regions with high confidence (<0.1% Irreproducibility rate) show significant (p < 0.05; Independent T-test) decrease in normalized TRF2 ChIP Seq reads in Super telomerase cells compared with wild type HT1080. The bars in above figures represent mean values of the replicates and error bars represent standard deviation. **D.TRF2 occupancy at multiple gene promoters was reduced with increase in telomere length in HT 1080-ST cells (D)** TRF2 occupancy at multiple promoter sites was reduced in HT 1080-ST cells in comparison with HT 1080 cells. **. E. Reduced occupancy of TRF2 on gene promoters lead to differential mRNA expression (E)** mRNA expression of 5 candidate genes in HT 1080 and HT 1080-ST cells **F. TRF2 silencing leads to increased** *p21* **promoter activity and expression. (F)**Transient TRF2 silencing using siRNA showed an increase in *p21* promoter activity and endogenous *p21* protein expression. **G**. **Telomere length dependent transcriptional regulation of p21 by TRF2 in HT 1080-ST cells (G)** *p21* promoter activity was relatively enhanced in HT 1080-ST than HT cells Endogenous *p21* (mRNA) and protein increased in HT 1080-ST relative to HT cells

In order to test the generality of our observations we decided to test the hypothesis in a completely different model of telomere elongation. An artificial telomere elongation model was reported by Wright's group where treatment of cells with ^18^ G-rich oligonucleotides (GTR) lead to telomere elongation. We replicated the model using immortalized MRC5 cells and found significant telomere elongation in cells sequentially fed with GTR for either 7 or 14 cycles (oligofed (OF) 7 or 14 cells, respectively in following text) (Figure 4A–B). We characterized the cells for expression of *hTERT, hTERC* and *TRF2* (Supplementary Figure 3B). Also, as expected, there was significantly more occupancy of TRF2 on telomeres in the oligofed cells (OF7, OF 14) (Figure 4C). We tested extra-telomeric TRF2 occupancy on the same sites that were validated in HT1080-ST cells earlier (Figure 3D) and found reduced TRF2 occupancy with increasing telomere length (Figure 4D). We then proceeded to check the effect of reduced TRF2 occupancy on the *p21* promoter on transcriptional output of the *p21* gene. We found that there was increase in promoter activity, mRNA expression and p21 protein levels in MRC5 OF7/OF14 cells (Figure 4 E), which is consistent with our observations with HT1080-ST cells.

**Figure 4.**
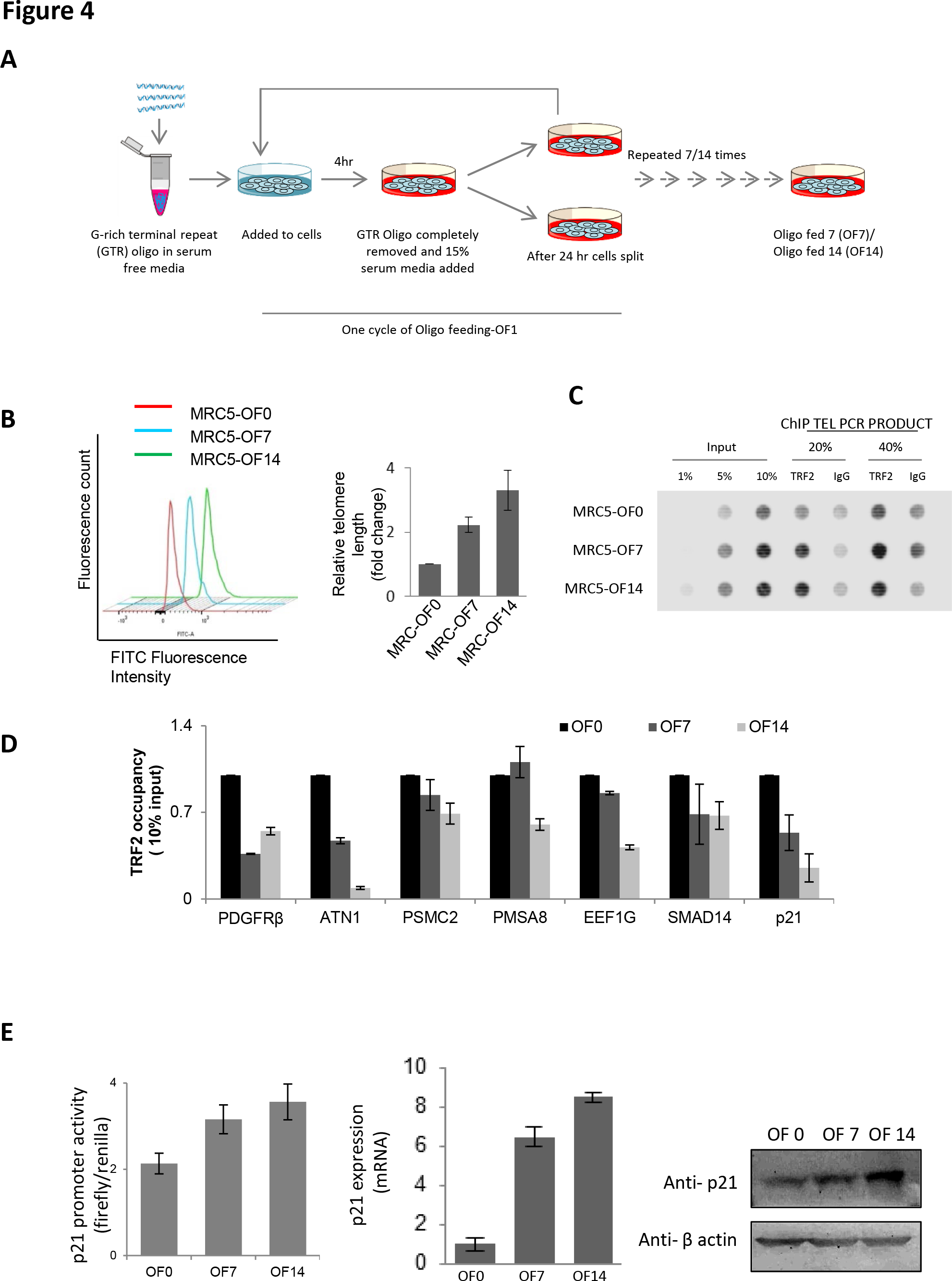
Reduced TRF2 occupancy at *p21* promoter in cells with artificially elongated telomeres leads to *p21* activation. **A-B.Telomere length elongation in MRC5 cells. (A)** Scheme showing steps followed in generating cells with artificially elongated telomere. Cells were treated with G-rich terminal repeats (GTR) [ (TTAGGG)4 100 mM] oligonucleotides in serum free media. After the treatment, GTR containing media was removed after 6 hrs. followed by media wash twice and cells were grown for 24 hrs. before splitting. Cycles were repeated to obtain cells with desired number of oligonucleotide feeding (OF). **(B)** Telomere elongation in oligo-fed MRC5 cells was confirmed by flow-FISH using FITC-tagged Telomere specific probe. **C. Telomere elongation in MRC5 cells show increased telomere occupancy of TRF2 (C)** ChIP with TRF2 antibody (or isotypic control) followed by PCR using telomere-specific primer in untreated, OF7 and OF14 cells showed that increase in telomere length resulted in increase in TRF2 occupancy at telomeres in OF cells (OF7, OF14) compared to untreated cells (OF0). **D.TRF2 occupancy at multiple gene promoters in MRC5 cells was reduced with increase in telomere length (D)** TRF2 occupancy at multiple promoter sites was reduced in OF7 and OF14 cells in comparison with untreated (OF 0) cells. **E. Telomere length dependent transcriptional regulation of *p21* by TRF2 in MRC5 oligo-fed cells (E)** Promoter activity, mRNA and protein expression of *p21* increased with increase in telomere length.

Extra-telomeric occupancy of TRF2 has been reported in previous ChIP-seq studies^5,11^. However, the functional significance for such genome wide extra-telomeric binding has remained largely unexplored. Results herein suggest that the telomeres can sequester TRF2 and control its extra-telomeric occupancy, and this change in extra-telomeric occupancy changes transcriptional landscape of cells. Our work presents the first evidence of transcriptional regulation of *p21* by TRF2 in a telomere length dependent manner. Given the importance of *p21* in cell cycle regulation and senescence, this observation is expected to lead to a better understanding of connection between cell cycle regulation and telomere length.

## Author Contributions

Following experiments were conducted and analyzed by: characterization of HT cells with altered telomere length and all experiments to support telomere length observations: AKM, SS; creation and characterization of MRC telomere length cells and all experiments to support telomere-length related observations in these cells: AKM, SS; all bioinformatics analyses including ChIP-seq/RNA-seq: PD; ChIP-seq, RNA-seq experiments and ChIP experiments: DS, AS, AK, AKM, SS; all microscopy experiments and flow cytometry experiments-SuSG, DS, TH, SDR, SSG, MK, AKM; design of experiments, manuscript preparation/editing-SC, AKM, PD, TH, DS, SS; conceptualization and overall execution of the research problem-SC.

## Acknowledgments

This work was funded by CSIR (GENCODE-A) and Wellcome Trust DBT India Alliance (500127/Z/09/Z). Research fellowship from CSIR to TH, DS, AKM, PD, AK and SDR is acknowledged. Research fellowship from ICMR to GP and AS is acknowledged. Research fellowship from Wellcome Trust DBT India Alliance to SS, SuSG and VS is acknowledged. We thank Peter Lansdorp for providing G4 specific 1H6 antibody and Joachim Lingner for providing HT1080-Supertelomerase cells as gifts. SC is senior research fellows of the Wellcome Trust/ DBT India Alliance.

---

**Supplementary Figure 1.**
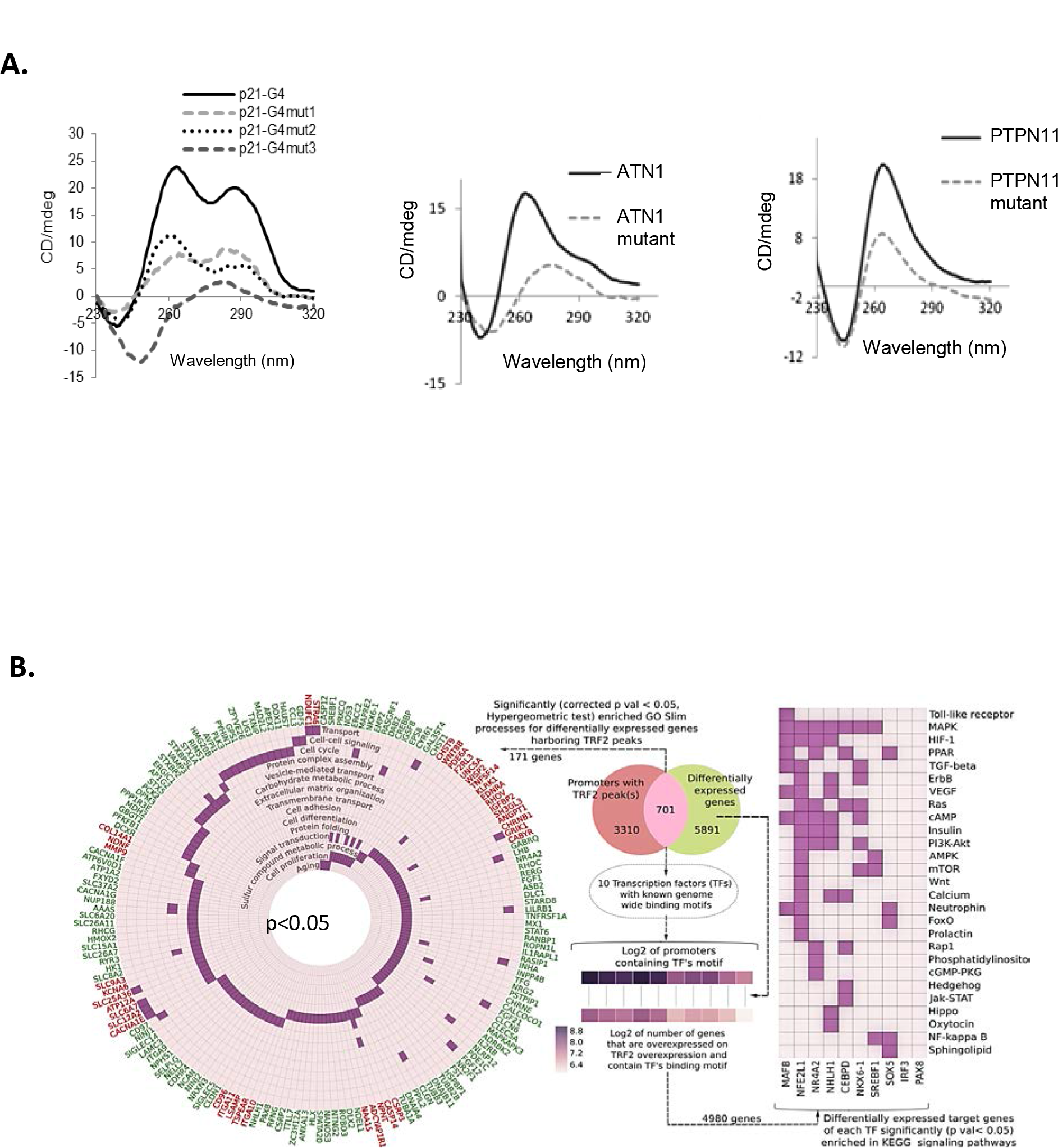
A. Promoter regions that harbor G4 motifs within TRF2 peaks form quadruplexes *in vitro*. Circular dichroism showing oligonucleotide constituting potential G4-motifs identified within TRF2 peak in 3 candidate genes (*p21, ATN1,PTPN11*) adopts G4 structure in solution and on substitution of bases required for G4 formation gives partial/complete disruption of the G4-motif under similar conditions. **B. TRF2 peaks showing differential gene expression:**701 differentially expressed genes had TRF2 peaks in promoter regions, 171 were enriched in GO Slim processes including signal transduction and cell cycle factors (p<0.05), and 50 (of 701) were transcription factors (Animal TFDB (Zhang et al., 2012)). For 10 of these transcription factors, predicted target binding sites (SwissRegulon (Pachkov et al., 2007) and ReMap (Griffon et al., 2015)) mapped within 10 kb of TSS of 4980 of the 5891 differentially expressed genes (entire altered transcriptome). These secondary TRF2-targets mapped to signaling pathways altered in cancer-4980 genes mapped to 27 pathways including Toll-like receptor, MAPK, TGF-beta and PI3K-Akt signaling (p<0.05; using KEGG API; Hyper-geometric test with Bonferroni correction).

**Supplementary Figure 2.**
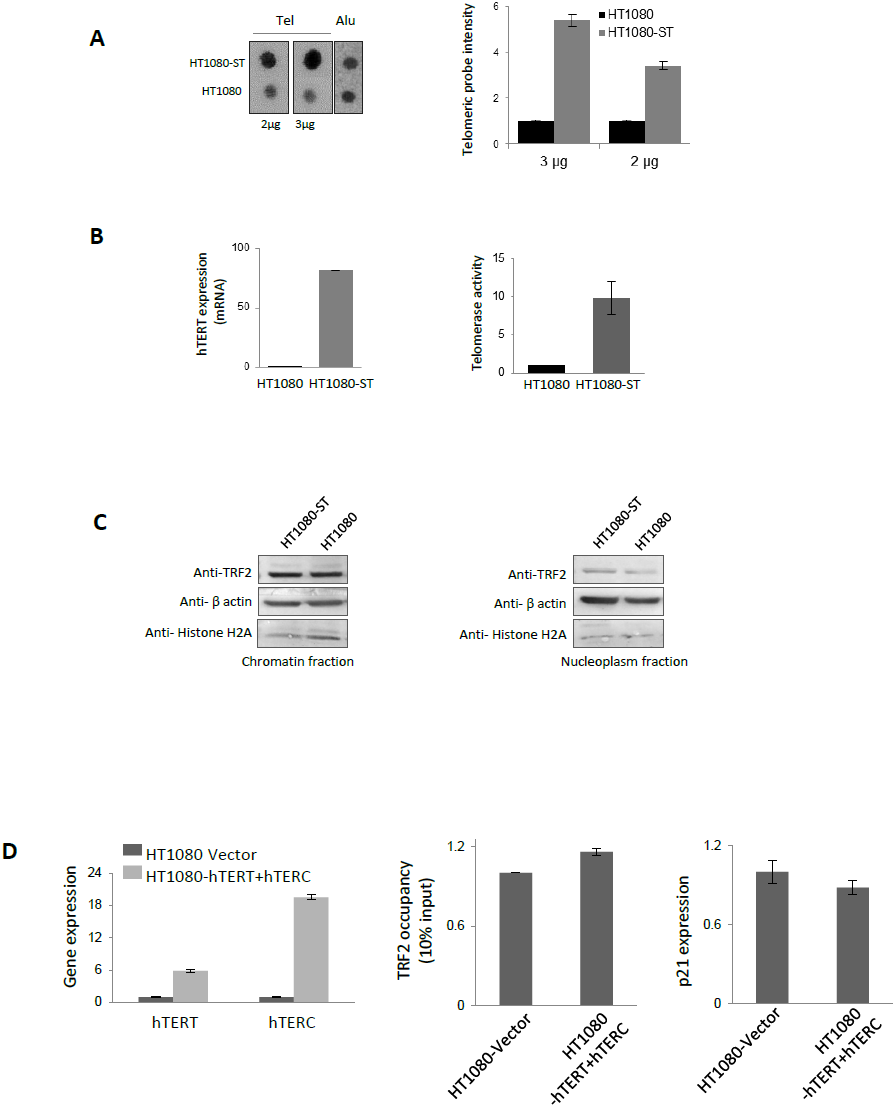
A. HT1080-ST cells harbor increased telomeric DNA. **(A)** Dot blot showing approximately 2 fold more telomeric probe intensity (normalized to Alu) for HT1080-ST cell as compared to HT1080 cells; **B. HT 1080-ST cells have higher *hTERT* expression and Telomerase activity (B)** HT1080-ST cells show enhanced *hTERT* levels (determined by qRT-PCR) and telomerase activity as determined by quantitative real-time TRAP(telomerase repeat amplification). **C. Both Nucleoplasm fraction and Chromatin bound TRF2 levels are similar in HT 1080 and HT 1080-ST cells (C)** Nuclear TRF2 levels were comparable in chromatin as well as nucleoplasm fraction in HT1080-ST and HT1080 cells. **D. Telomerase over expression does not cause change in TRF2 occupancy and p21 regulation (D)** Upon transient overexpression of hTERT and hTERC in HT1080 cells,there was no significant change in TRF2 occupancy at the p21 promoter and p21 expression.

**Supplementary Figure 3.**
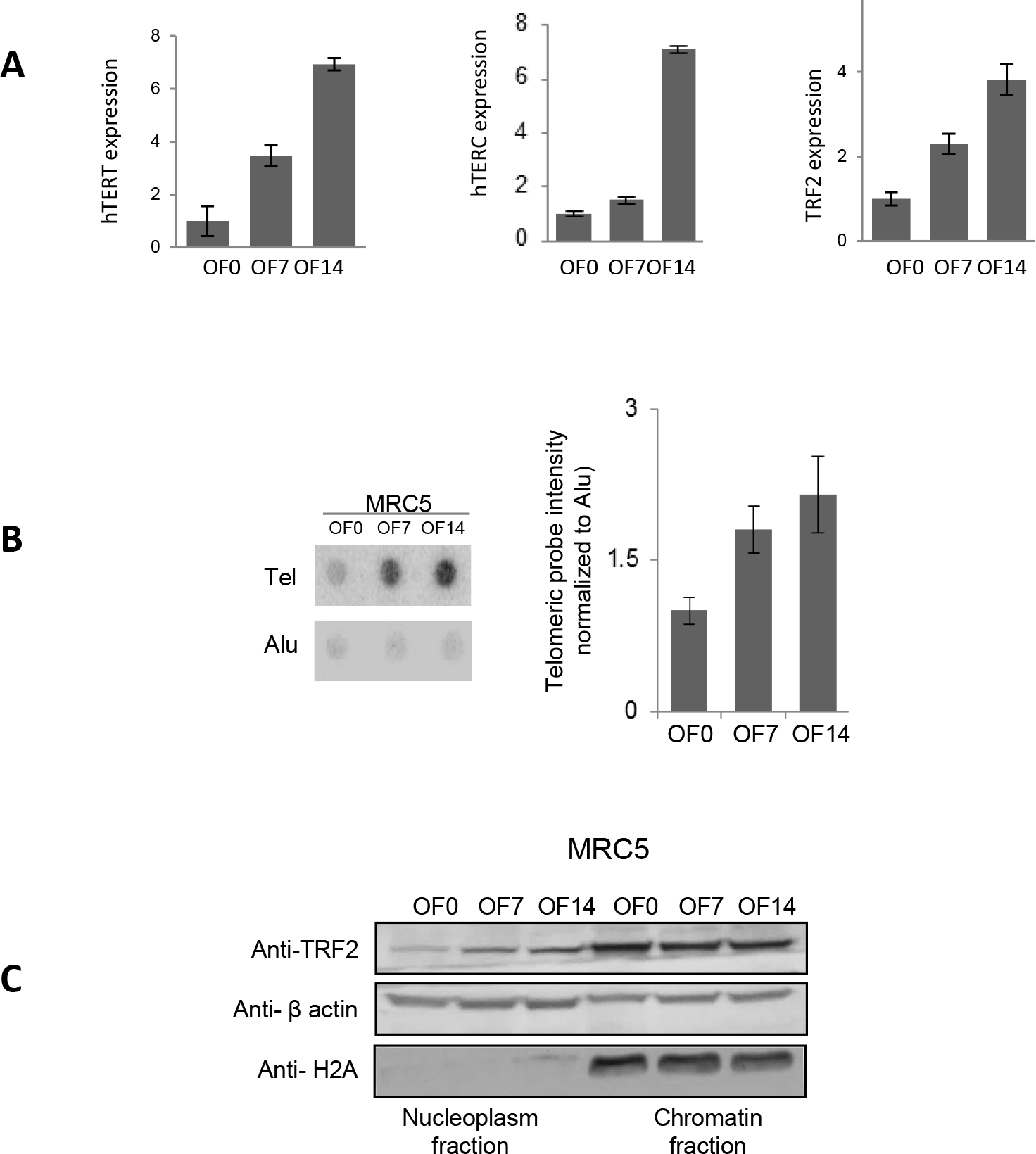
A. MRC5 oligo fed cells have higher *hTERT* and expression (B) MRC5 oligofed cells (OF7,OF14) show enhanced *hTERT* and levels (determined by qRT-PCR) and TRF2 mRNA levels are also enhanced in oligofed cells (OF7,OF14). **B**.**MRC5 oligofed cells have increased telomeric DNA. (B)** Dot blot showing higher Telomeric probe signal (normalized to Alu) in MRC5 oligofed cells (OF7,OF14) **C. Chromatin bound TRF2 levels are similar in MRC5 oligofed cells in comparison to untreated MRC5 cells(C)** Nuclear TRF2 levels were comparable in chromatin fraction in MRC5 oligofed cells (OF7,OF14) with untreated cells (MRC5 OF 0).

## Online Methods

### Cell lines, media and culture conditions

HT1080 fibrosarcoma cell line was purchased from the NCCS, Pune (India). Immortalized MRC5 cells were received as a gift from NII, New Delhi (India). MDA-MB-231 cells were received as a gift from Mayo Clinic, Minnesota (USA). HT1080 and MRC5 cells were maintained in Modified Eagle's medium (MEM) supplemented with 10% Fetal Bovine Serum (FBS). MDA-MB-231 and HCT116 cells were maintained in Dulbecco's Modified Eagle's medium (DMEM) supplemented with 10% FBS. All cultures were grown in incubators maintained at 37°C with 5% CO2.MRC 5 oligofed cells were made by periodic treatment of oligofed cells with GTR (G rich Terminal repeat) sequences [(TTAGGG)4 100 mM] as explained in schematic Figure 4A (main manuscript)

### Flow-FISH

The Flow-FISH assay for telomere length detection was performed using DAKO Telomere PNA Kit/FITC codeK5327. For each sample, 3 million cells were counted and divided into three parts: pelleted down and washed with 1X PBS (two times). One part was fixed with chilled 75% ethanol and stored at -20°C. The other two parts were resuspended in hybridization solution (with and without FITC labeled PNA probe). The cells were incubated at 82°Cfor 10 mins and stored in dark at RT for 12 hrs. The hybridization buffer was washed off with 1X PBS and the cells were incubated with Propidium Iodide (PI) solution provided with the kit at RT for 2 hrs. The cells diluted with PBS and flow cytometry was performed using BD ARIA III flow cytometer. The ethanol fixed cells were incubated with PI solution and flow cytometry was performed using BD ARIA III flow cytometer for cell cycle analysis. All downstream analysis of data was done using recommendations of the kit.

### Immunofluorescence microscopy

Cells were grown on cover slips and at 100% confluence were fixed with 4% Paraformaldehyde by incubating for 10 min at RT. Cells were permeabilized with 0.5% Triton™ X-100 and treated with blocking solution (3% BSA in PBS) for 30 min at RT. After one PBS wash cells were treated with anti-TRF2 antibody (1:100) and anti-G4 1H6 antibody (1:150, a kind gift from Dr. Peter M. Lansdorp, BC Cancer Agency, Vancouver, Canada, for Fig. 2D; incubated overnight at 4°C. Next day cells were washed alternately with PBS and PBST three times and probed with Alexa Fluor^®^ 488 and Alexa Fluor^®^ 594 which required an incubation for 2 hr at RT. Cells were washed again alternately with PBS and PBST three times and mounted with Prolong^®^ Gold anti-fade reagent with DAPI. Images were taken as Maximum Intensity Projections on Leica TCS-SP8 confocal microscope. TRF2-G4 co-localization (Fig. 2D) was calculated using Volocity 6.3 software as Global Pearson's Correlation.

### Library preparation for ChIP-Seq and RNA-Seq

TRF2-bound DNA from HT-1080 cells was quantified, and 20 ng from each sample was taken for end repair using Illumina Tru-Seq sample preparation kit. Samples were purified using PCR purification kit (Qiagen, Germany). Thereafter ‘A’ base was added to the samples 3′-end using Illumina sample preparation kit. After the end of the reaction, samples were again purified by PCR purification kit (Qiagen). Then flow-cell primer specific adapters were ligated to the ChIP DNA fragments and samples were further purified by MinElute columns. Size selection was done after adapter ligation using 2% agarose gel. Gel extraction columns (Qiagen) were used to purify DNA fragments ranging between 150 and 350 bases. These eluted samples were then purified using MinElute columns and these samples were then amplified for 18 cycles to enrich adapter-ligated DNA fragments. After PCR purification and elution the DNA was quantified using Picogreen method, and then 7 pico moles of each sample was sequenced on GAIIx or HiSeq2500 (Illumina, USA) according to manufacturer's protocol.

For the mRNA-Seq sample preparation, the Illumina standard kit was used according to the manufacturer's protocol. Briefly, 10μg of each total RNA sample was used for polyA mRNA selection using streptavidin-coated magnetic beads, followed by thermal mRNA fragmentation. The fragmented mRNA was subjected to cDNA synthesis using reverse transcriptase (SuperScript II) and random primers. The cDNA was further converted into double stranded cDNA and, after an end repair process (Klenow fragment, T4 polynucleotide kinase and T4 polymerase), was finally ligated to Illumina paired end (PE) adaptors. Size selection was performed using a 2% agarose gel, generating cDNA libraries ranging in size from 200-250 bp. Finally, the libraries were enriched using 15 cycles of PCR and purified by the QIAquick PCR purification kit (Qiagen). The enriched libraries were diluted with Elution Buffer to a final concentration of 10 nM. Each library was run at a concentration of 7 pM on Hi-seq lane using 76 bp pair end sequencing.

### Analysis of high throughput sequencing data

ChIP-Seq data was aligned to human reference genome (hg19) using Bowtie 2.1 short read aligner. Peaks were called using MACS 1.4.2 (shift size: 125, p values cut-off: 10e-5, FDR cut-off: 5%). Common peaks were identified as those peaks whose coordinates intersected using BEDTool's intersect sub-command. For downstream analysis common peaks were used (peak coordinates were slopped so that the start coordinate is the minimum of replicate 1 and replicate 2 and end coordinate is the maximum in replicate 1 and replicate 2).RNA-Seq reads were aligned to human reference genome (hg19) using TopHat v2.0.9. Gene wise quantification of expression was performed using Cufflinks v2.2. Genes were labelled as up-regulated and down-regulated with fold change cut-off of 1.5 and 1/1.5 respectively. For comparison of HT1080 and HT1080-ST we compared TRF-ChIP RPM values from two samples (raw reads per million sequenced reads) for 1226 genomic regions (these genomic regions are peaks called using MACS that had Irreproducibility Discovery Rate (IDR) less than 0.1%).

### Integrated analysis of sequencing data with PG4 motifs

PG4 motifs were identified from human reference genome (hg19) using custom Python scripts. The regular expression used to map G3L1-x motif used was: (?:G{3}[ATGCN]—1,x}){3}G{3} and was implemented using 're’ module in Python. PG4 motifs were mapped to ±10kb of transcription start sites (obtained from UCSC RefSeq genes) using Pybedtools. To test for significance of PG4 motifs in common peaks, each peak was shuffled 1000 times followed by search for PG4 motif in the shuffled peak followed by test of significance using Pearson's chi-squared test as implemented in Python's Scipy package. To further test for association of motif with TRF2 peaks. Distance of nearest PG4 motif from each TRF2 was calculated using BED tools closest sub-command. Simultaneously, a shuffled control dataset was generated so that there is a random genomic coordinate corresponding to length of each peak. Both datasets were filtered to include to include peaks/shuffled coordinates present in 10kb proximity of the TSS. Contingency table was constructed as shown below.

|  |  |
| --- | --- |
| $\Sigma C_{\text{TRF2}}$ | $\Sigma C_{\text{shuffle}}$ |
| $\Sigma R_{\text{TRF2}}$ | $\Sigma R_{\text{shuffle}}$ |

Where, ∑C represents sum of number of peaks/shuffle coordinates having PG4 in three central bins(i.e. in proximity to TSS), ∑R represents sum of number of peaks/shuffle coordinates having PG4 in rest bins. Fisher's exact test was performed on this contingency table using Scipy package in Python. Further, reads were binned in 10bp window and mapped to +−10kb of TSS of RefSeq genes using python package deepTools. Thereafter, differentially expressed genes with a common peak in replicates were plotted in Figure 1(i) in bins of 100bp using python packages MetaSeqand Pybedtools.

### ChIP (Chromatin Immunoprecipitation)

ChIP assays were performed as per protocol provided by Upstate Biotechnology with modifications as suggested in Fast ChIP protocol. ChIP assays were performed using anti-TRF2 antibody(Novus Biologicals NB110-57130). Anti-Rabbit IgG was used for isotype control in all cell lines. Briefly, cells were fixed with ∼1% formaldehyde for 10min and lysed. Chromatin was sheared to an average size of ∼500-1000 bp using Biorupter (Diagenode). 10% of sonicated fraction was processed as input using phenol-chloroform and ethanol precipitation. ChIP was performed using 1:100 dilution (v/v) of the respective antibody incubated overnight at 4°C. Immune complexes were collected using herring sperm DNA-saturated Magnetic Dyna beads and washed extensively. Phenol-Chloroform-Isoamylalcohol was used to extract DNA from immunoprecipitated fraction. ChIP DNA was further validated by using either semi-quantitative PCR or qRT-PCR method.

### Circular dichroism

The circular dichroism (CD) spectra were recorded on a Jasco-810 Spectropolarimeter equipped with a Peltier temperature controller. Experiments were carried out using a 1mm path-length cuvette over a wave length range of 200-320 nm. 5uM oligos were diluted in KCl buffer (10mM HEPES and 100mM KCl, pH 7.4) and denatured by heating to 95°C for 5 min and slowly cooled to 25°C for overnight. The CD spectra reported here are representations of three averaged scans taken at 25 °C and are baseline corrected for signal contributions due to the buffer.

### TRF2 silencing

HT1080 cells were transfected with TRF2 siRNA oligonucleotides synthesized from Dharmacon using lipofectamine 2000 (Invitrogen) transfection reagent according to manufacturer's instructions. Silencing was checked after 48 hr of transfection. Pooled SCR siRNA was used as control.

### Luciferase assay

Minimal promoter region of p21 harboring TRF2 binding peak were cloned into pGL3 vector between Kpn1 and HindIII restriction sites. Promoter region was amplified by using 5‘-GACTGGGCATGTCTGG-3’ as forward primer and 5‘-CTCTCACCTCCTCTGAGTG-3’ as reverse primer from genomic DNA of human WBC. Firstly, this amplified region was ligated into TA vector and then subcloned into pGL3 vector by Kpn1 and HindIII restriction enzyme and sequence was verified. Plasmid (pGL4.73) containing a CMV promoter driving Renilla luciferase was cotransfected as transfection control for normalization. After 48h, cells were harvested and luciferase activities of cell lysate were recorded by using a dual-luciferase reporter assay system (Promega).

### Real time PCR

Total RNA was isolated using TRIzol^®^ Reagent (Invitrogen, Life Technologies) according to manufacturer's instructions. A relative transcript expression level for genes was measured by quantitative real-time PCR using a SYBR Green based method. Average fold changes was calculated by difference in threshold cycles (Ct) between test and control samples. *GAPDH* gene was used as internal control for normalizing the cDNA concentration of each sample.

### Dot blot analysis

For dot blot analysis, Genomic/ ChIP DNA was denatured at 95°C and dot blotted on N+ hybond membrane (Amersham) in pre-wetted in 2X SSC buffer. The DNA was UV cross-linked. Membranes were pre-hybridized in Rapid-Hyb buffer (Amersham) for 30 mins at 37°C. Following this, hybridization with a 24-bp radio-labeled telomeric probe (AATCCC)_4_ or 418-bp radio-labelled ALU probe as performed for 4h at 42°C and 65°C respectively; and membranes washed with 2X SSC and 0.2X SSC + 0.1% SDS twice at hybridization temperature before exposing overnight on phosphoimager imaging plate. All data were scanned using Bio-Rad Personal Molecular Imager. Data was processed and quantified using Image J image analysis software.

### Chromatin and nucleoplasm fractionation assay

Chromatin fractionation assay was carried out as described (Lou et al., 2006). Briefly, the cells were lysed in buffer I (50 mM HEPES, pH 7.5, 150 mMNaCl, 1 mM EDTA, 0.05% NP40, and protease and phosphatase inhibitors) for 5 minutes on ice. Cell lysate was centrifuged at 3500 rpm for 5 minutes at 4°C. Supernatant was collected as nucleoplasm fraction (fraction I). Further the precipitate was washed once with buffer I and then resuspended in buffer II (50 mMTris-HCl, pH 7.5, 150 mMNaCl, 1% NP40, 0.5% sodium deoxycholate, 0.1% SDS and protease and phosphatase inhibitors) and incubated in ice for 20 minutes. Next re-suspended fraction was centrifuged at 14000 rpm for 20 minutes at 4°C and supernatant was collected as the chromatin-enriched fraction (fraction II).

### Western blotting

For western blot analysis, protein lysates were prepared by suspending cell pellets in 1X cell culture lysis buffer (promega). Protein was separated using 12% SDS-PAGE and transferred to polyvinylidenedifluoride membranes (Immobilon FL, Millipore). After blocking the membrane was incubated with primary antibodies-anti-TRF2 antibody (Novus Biological), anti-p21 antibody (Cell signaling technology) and anti-P-actin/anti-GAPDH antibody (Sigma). Secondary antibodies, anti-mouse and anti-rabbit alkaline phosphatase conjugates were from Sigma. The blot was finally developed by using Thermo Scientific Pierce NBT/BCIP developing reagents.

